# Temporal perturbation of Erk dynamics reveals network architecture of FGF2-MAPK signaling

**DOI:** 10.1101/629287

**Authors:** Yannick Blum, Jan Mikelson, Maciej Dobrzyński, Hyunryul Ryu, Marc-Antoine Jacques, Noo Li Jeon, Mustafa Khammash, Olivier Pertz

## Abstract

Stimulation of PC-12 cells with epidermal (EGF) versus nerve (NGF) growth factors (GFs) biases the distribution between transient and sustained single-cell ERK activity states, and between proliferation and differentiation fates within a cell population. We report that fibroblast GF (FGF2) evokes a distinct behavior that consists of a gradually changing population distribution of transient/sustained ERK signaling states in response to increasing inputs in a dose response. Temporally-controlled GF perturbations of MAPK signaling dynamics applied using microfluidics reveals that this wider mix of ERK states emerges through the combination of an intracellular feedback, and competition of FGF2 binding to FGF receptors (FGFR) and heparan-sulfate proteoglycans (HSPGs) co-receptors. We show that the latter experimental modality is instructive for model selection using a Bayesian parameter inference. Our results provide novel insights into how different receptor tyrosine kinase (RTK) systems differentially wire the MAPK network to fine tune fate decisions at the cell population-level.

Microfluidics, Erk Signaling Dynamics, Mechanistic Modelling, Parameter Estimation, Cell Fate Determination

## Introduction

Signaling dynamics, rather than steady states, have been shown to control cell fate responses (Levine et al., 2013). For multiple systems including receptor tyrosine kinase signaling (RTK), signaling heterogeneity can explain the fate variability observed within a cell population (Chen et al., 2012; Cohen-Saidon et al., 2009). Both, biological noise extrinsic to individual cells and intrinsic variability within signaling networks shape the cell fate. It has been proposed that the dynamic nature of signal transduction enables accurate information transmission in the presence of noise (Wollman, 2018). Measuring single-cell signaling dynamics is therefore key to understanding how cellular responses correlate with specific cell fate decisions.

The extracellular-regulated kinase (ERK) is a key regulator of fates such as proliferation and differentiation. It functions within a mitogen-activated protein kinase (MAPK) signaling pathway in which growth factor (GF) receptors activate a membrane-resident Ras GTPase that subsequently triggers a MAPK cascade leading to ERK activation (Avraham and Yarden, 2011). Rat adrenal pheochromocytoma PC-12 cells have been widely used as a model system to study the regulation of specific cell fates by MAPK signaling (Marshall, 1995). Stimulation with EGF or NGF leads to population-averaged transient or sustained ERK states, which specifically trigger proliferation or differentiation. Thus, ERK signal duration has been proposed as a key determinant of cell fate (Marshall, 1995; Santos et al., 2007). These distinct ERK states result from GF-dependent control of the MAPK network (Santos et al., 2007), with negative and positive feedback producing all-or-none adaptive or bistable outputs, respectively (Avraham and Yarden, 2011; Santos et al., 2007; Xiong and Ferrell, 2003). More recently, single-cell assays have indicated that EGF/NGF induce heterogeneous dynamic signaling states across a cell population (Ryu et al., 2015). While EGF leads to transient ERK activity responses, NGF induces transient or sustained responses in an isogenic population due to variability in expression of signaling components and receptor-dependent modulation of the negative and positive feedback loops. This might explain how NGF can induce a heterogeneous mix of differentiating and proliferating cells (Chen et al., 2012). Further support that dynamic ERK signaling states control fate decisions stems from model-based prediction of dynamic GF stimulation schemes that induce synthetic ERK activity patterns that determine fate decision independently of GF identity (Ryu et al., 2015).

FGF2 activates ERK through FGF receptors (FGFRs) and regulates processes such as angiogenesis, wound healing and development (Ornitz and Itoh, 2015). Upon FGF2 stimulation, FGFR dimerizes, autophosphorylates, recruits adaptors, and activates the Ras/RAF/MEK/ERK cascade (Ornitz and Itoh, 2015). In PC-12 cells, FGF2 induces sustained ERK activity, which correlates with differentiation (Qui and Green, 1992). FGF-FGFR interactions are further regulated by a heparan sulfate proteoglycan co-receptor (HSPGs) (Matsuo and Kimura-Yoshida, 2013; Ornitz, 2000). FGF2 initially binds to HSPGs through a high affinity interaction, followed by a 2^nd^ lower affinity interaction leading to a HSPG/FGF2/FGFR trimeric complex. The latter subsequently dimerizes to a dimer of trimers complex that can autophosphorylate and signal downstream (Ornitz and Itoh, 2015). In marked contrast with signaling systems that exhibit sigmoidal dose responses, FGF2 elicits biphasic signaling and cell fate output responses. For example, bell-shaped neuronal differentiation (Williams et al., 1994), or cell proliferation fate outputs (Zhu et al., 2010) are observed in FGF2 dose response challenges in different cell systems. This correlates with biphasic ERK activity outputs (Kanodia et al., 2014; Zhu et al., 2010). The ability of FGF2 to induce biphasic responses has been proposed to emerge from competition of FGF2 binding to HSPGs and the FGFR (Kanodia et al., 2014).

Here, we explore how the FGF2/FGFR controls ERK activity dynamics at the single-cell level in PC-12 cells. We find that FGF2 induces a mix of dynamic ERK states that are distinct from those of EGF/NGF, with increasing FGF2 input gradually modulating the distribution of transient/sustained ERK states. Our data together with mathematical modelling show that the FGF2-dependent MAPK signaling network underlying these responses consists of an extracellular FGF2/FGFR/HSPG interaction layer coupled to an intracellular MAPK network layer with a simple negative feedback. We conclude that EGF, NGF, FGF2 wire the MAPK differently to induce distinct population distributions of ERK states that fine-tune fate decisions at the cell population level.

## Results

### FGF2 induces dynamic signaling states distinct from those induced by EGF/NGF

EGF/NGF-triggered ERK activity responses have been widely studied in PC-12 cells. However, single-cell studies have revealed a much higher signaling complexity than previously anticipated (Ryu et al., 2015). Here, we asked if FGF2 potentially induces ERK activity dynamics within a cell population that are distinct from those of EGF/NGF. To study FGF2 signaling at the single-cell level, we used a PC-12 cell line stably expressing EKAR2G, a fluorescence resonance energy transfer (FRET)-based biosensor for endogenous, cytosolic ERK activity. EKAR2G has been extensively validated elsewhere (Fritz et al., 2013; Harvey et al., 2008). To extract single-cell temporal ERK activity patterns (scTEPs), we used a CellProfiler-based (Kamentsky et al., 2011) image analysis pipeline for segmentation and tracking of single cells, and for computation of a per-cell average FRET biosensor ratio. We used a computer-programmable microfluidic device to temporally perturb cells using GF pulses (Fig 1A).

**Figure 1.**
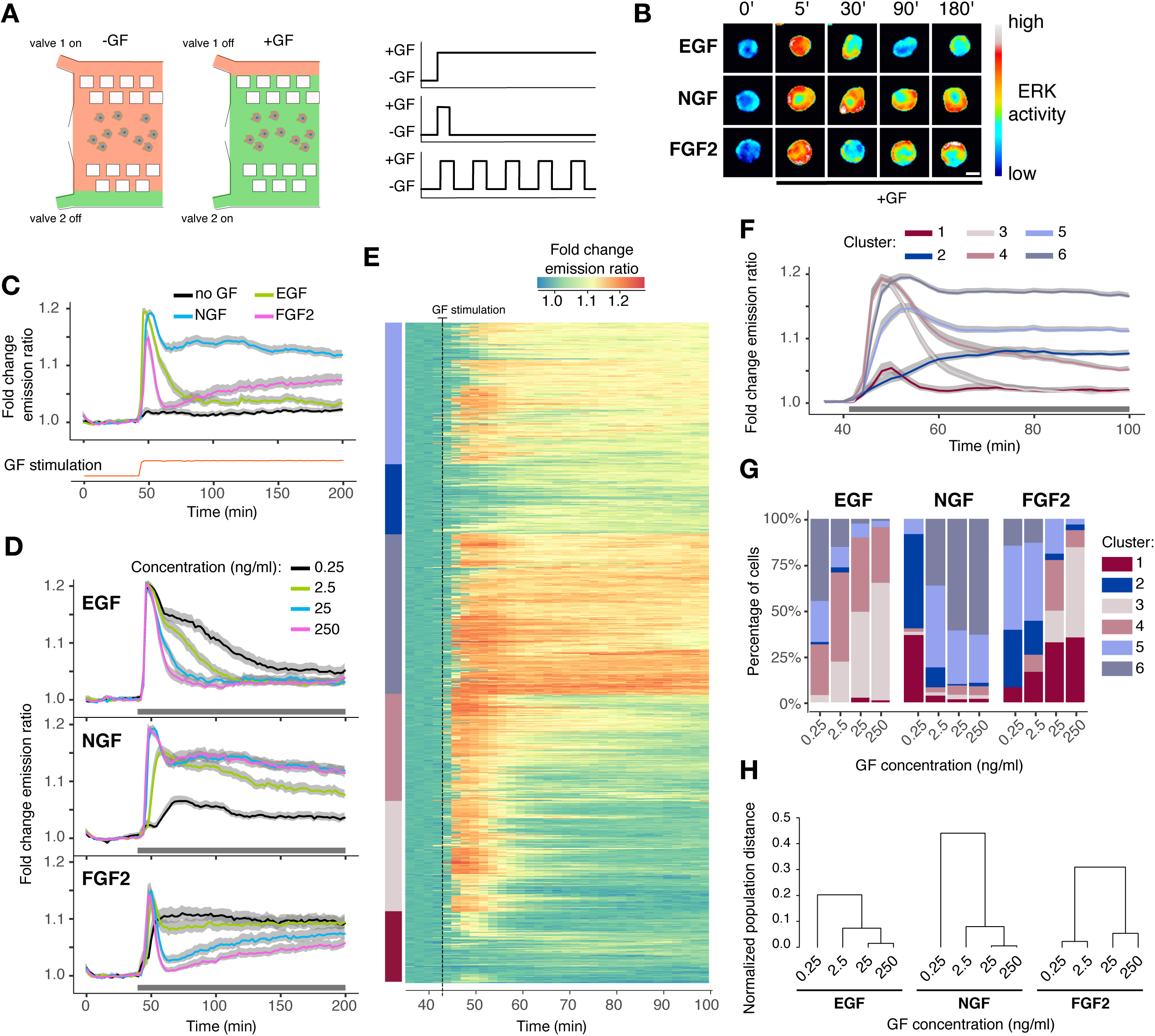
FGF2 induces different dynamic ERK activity signaling states than EGF/NGF. (A) Flow-based, microfluidic device for temporal GF delivery. Computer-controlled, pressure pump enables mixing of medium and GFs to deliver GF pulses in cells cultured in the microfluidic device. Right panel illustrates typical GF stimulation patterns. (B) Representative EKAR2G ratio images of cells treated with 25 ng/ml EGF, NGF and FGF2. Ratio images are color-coded so that warm/cold colors represent high/low ERK activation levels. Scale bar = 50 ⌈m. (C,D) Population averages of ERK activity dynamics in response to stimulation with 25 ng/ml (C), or with a dose-response challenge using 0.25, 2.5, 25, 250 ng/ml EGF, NGF, FGF2 (D). Single-cell time series were normalized to their own means before GF stimulation, t = [0, 40]. Red curve at the bottom of panel C indicates GF stimulation profile measured simultaneously using an Alexa-546-labeled dextran. N=[48, 120] cells per GF concentration. ERK dynamics measured at 2’ intervals. (E) Hierarchical clustering of pooled (N = 983) single-cell time series from panel D. To focus on relevant ERK dynamics, we trimmed x-axis to t = [36, 100] min. Each row of the heatmap corresponds to a time series of a single cell. We used Dynamic Time Warping as a distance measure and Ward’s minimum variance linkage method for building the dendrogram, which was then cut to distinguish 6 clusters that are color-coded on the left. (F) Average ERK activity across 6 clusters identified in panel E, color-coded as in (E). (G) Distribution of ERK activity trajectories across 6 clusters from panels E, F in response to different GF dosages. (H) Separability between populations of single-cell trajectories calculated as normalized area under curve of Jeffries-Matusita distance along time (Methods, Fig. S2B). The dendrogram was created using the complete-linkage method. In panels C, D, and F: grey band indicates 95% CI for the mean. In panels D and F: black horizontal bar indicates GF stimulation.

First, we stimulated cells with a typical EGF/NGF/FGF2 concentration of 25 ng/ml (Fig 1B,C). We used a fluorescent dextran for quality control of the GF delivery by the microfluidic device (Fig 1C, lower red trace). As expected, when we evaluated population-averaged temporal ERK activity patterns (paTEPs), EGF led to transient ERK activity, while NGF induced a peak followed by sustained ERK activity with an amplitude lower than the peak. In contrast, FGF2 led to a transient ERK peak that was sharper than the one evoked by EGF. After this fast adaptation, ERK activity gradually increases over time. Since increasing FGF2 concentration induces biphasic fate and ERK activity in a variety of cell systems (Kanodia et al., 2014; Zhu et al., 2010), we also tested such increase in our system. We stimulated PC-12 cells with EGF/NGF/FGF2 concentration in a 0.25 – 250 ng/ml range, as in previous works (Kanodia et al., 2014; Zhu et al., 2010) (Fig 1D). On average, all EGF concentrations triggered an initial ERK peak with identical amplitude but with faster adaptation at higher GF concentrations. In contrast, 0.25 ng/ml NGF only induced moderate ERK activity without an initial ERK activity peak. 2.5 ng/ml NGF led to sustained ERK activity after a small initial peak. 25 and 250 ng/ml led to almost indistinguishable profiles of an ERK activity peak followed by sustained ERK activity. FGF2 stimulation led to different paTEPs than both EGF and NGF. Indeed 0.25 ng/ml FGF2 led to sustained ERK activity without a robust initial peak, whereas 2.5, 25 and 250 ng/ml FGF2 led to a clearly defined initial ERK transient. At 25 and 250 ng/ml FGF2, after the initial transient, we again observed slow ERK activity recovery. The previously described biphasic ERK activity output is evident when we consider a time point after the initial ERK activity peak (Kanodia et al., 2014; Zhu et al., 2010).

### FGF2 dose response leads to an ERK activity population distribution that is wider than those associated with EGF and NGF

Heterogeneous single-cell dynamic signaling states were evident when scTEPs were overlaid over paTEPs (Fig EV1A). To examine this heterogeneity, we pooled all trajectories (EGF, NGF, FGF2 – 4 concentrations) using a time interval ranging from shortly before GF stimulation to 60’ after stimulation (Fig 1E). We then subjected this dataset to hierarchical clustering with dynamic time-warping (DTW) distance measure to extract classes of scTEPs based on their shape. Visual inspection of the resulting clusters led us to empirically choose different time intervals/datasets to highlight specific features of dynamic ERK activity states. In Figure 1F we identified 6 scTEP clusters (Fig 1F). ScTEPs/paTEPs for each cluster are shown in Figure EV1B. Even though all clusters differ in amplitude, we can recognize adaptive behavior in clusters 1, 3, 4, and sustained activation in clusters 2, 5, 6. We then computed the population distribution of these representative scTEPs across all experimental conditions (Fig 1G). Low-amplitude adaptive and sustained ERK activities (clusters 1 and 2) were largely absent from responses to all EGF stimulations, indicating robust ERK signaling. With the increase of EGF, an adaptive cluster 3 replaced sustained clusters 5 and 6. In contrast, the lowest NGF dose induced a mix of low-amplitude adaptive and sustained responses. High NGF concentrations induced high-amplitude sustained clusters 5 and 6 with a decreasing contribution of intermediate responses. We observed a wider mix of cluster distribution for FGF2 dose response. 0.25 ng/ml FGF2 led to a mix of low and high amplitude sustained responses. 2.5 ng/ml FGF2 decreased sustained responses in favor of cluster 4 (high amplitude, intermediate adaptation). Then, with an increased FGF2 dosage the distribution shifted to strongly adaptive responses with low and high amplitudes.

Intrigued by the fact that FGF2 induces slow ERK activity recovery after the 1st peak at 25 and 250 ng/ml in population-averaged measurements (Fig 1D), we tested if this was also the case at the single-cell level. For that purpose, we individually clustered the EGF, NGF, FGF2 dose response experiments on a time ranging from shortly before GF addition to 160’ after stimulation (Fig EV1C). For FGF2, this again identified scTEPs that displayed a robust 1st ERK activity peak followed by different levels of adaptation (clusters 1-3), or sustained ERK activity. Importantly, the 3 adaptive clusters, that were present at high FGF2 concentrations, displayed slow ERK activity recovery after adaptation. This specific phenomenon is not present in EGF/NGF dose responses.

To independently assess that FGF2 evokes a distinct and wider mix of scTEPs than EGF/NGF in a dose response, we both applied PCA decomposition and calculated accumulated pairwise distances of the response distribution at different timepoints (Figs 1H, (Fig EV2)). Both approaches showed that scTEPs population distributions for different GF dosages are more separated for FGF2. We discuss it further in Appendix text 1 and Materials and Methods.

### Decoding FGF2/MAPK signaling network properties by temporal perturbation of ERK dynamics

We then sought to identify the signaling network structure that explains how the FGFR/MAPK network evoke ERK states different from those evoked by EGF/NGF. For that purpose, we dynamically perturbed cells by delivering single or multiple GF pulses of different lengths and concentrations using our microfluidic device (Fig 1A). This approach captures salient features of the MAPK network not accessible with sustained GF stimulation, and in many cases, induces population-homogeneous signaling states that are simpler to interpret (Ryu et al., 2015). We stimulated PC-12 cells with pulses of 3’,10’ and 60’ with the four concentrations of each GF used previously. We plotted the paTEPs (Fig 2) and used hierarchical clustering to extract representative dynamic patterns for each GF pulse pattern (Fig EV3).

**Figure 2.**
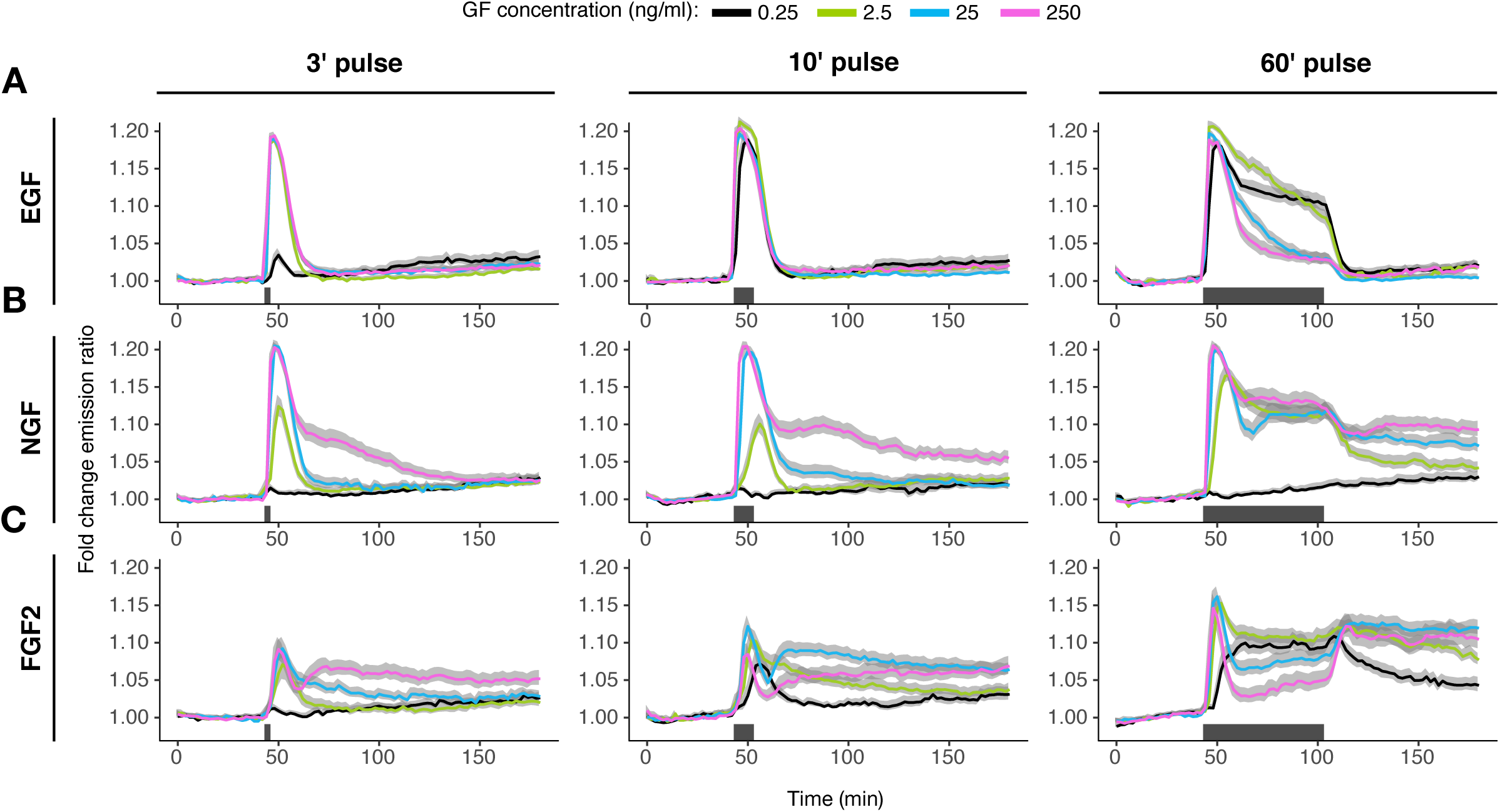
ERK activity dynamics in response to single-pulse stimulation. (A,B,C) Population average ERK activity dynamics in response to 3’, 10’, 60’ EGF (A), NGF (B), FGF2 (C) single pulse stimulation. Single-cell time series were normalized to their own means before the GF stimulation, t = [0, 40]. Solid lines – population mean, N=[39, 166]; grey bands – 95% CI for the mean; black horizontal bars – duration of GF stimulation.

The pulsed EGF/NGF dose responses were consistent with our previous observations (Ryu et al., 2015). PaTEPs exhibited a full-amplitude initial ERK activity peak followed by robust adaptation for all EGF concentrations for 3’ or 10’ pulse, except for a 3’ 0.25 ng/ml EGF pulse (Fig 2A) where the peak was less pronounced. The 60’ EGF pulse revealed distinct adaptation kinetics after the initial ERK activity peak with faster adaptation at higher EGF dose. As observed in sustained stimulation, full adaptation occurred concomitantly with EGF washout. Clustering of scTEPs revealed adaptive responses across the EGF doses and pulsing schemes (Fig EV3A). In the case of NGF, the 0.25 ng/ml concentration did not yield ERK activation across any pulsing scheme (Fig 3B). Above this concentration, high NGF input (achieved by increasing dose and/or pulse duration) gradually shifted the paTEPs from transient to more sustained profile. Clustering of scTEPs revealed a mix of transient and sustained responses, with sustained clusters contributing more at high NGF inputs (Fig EV3B).

**Figure 3.**
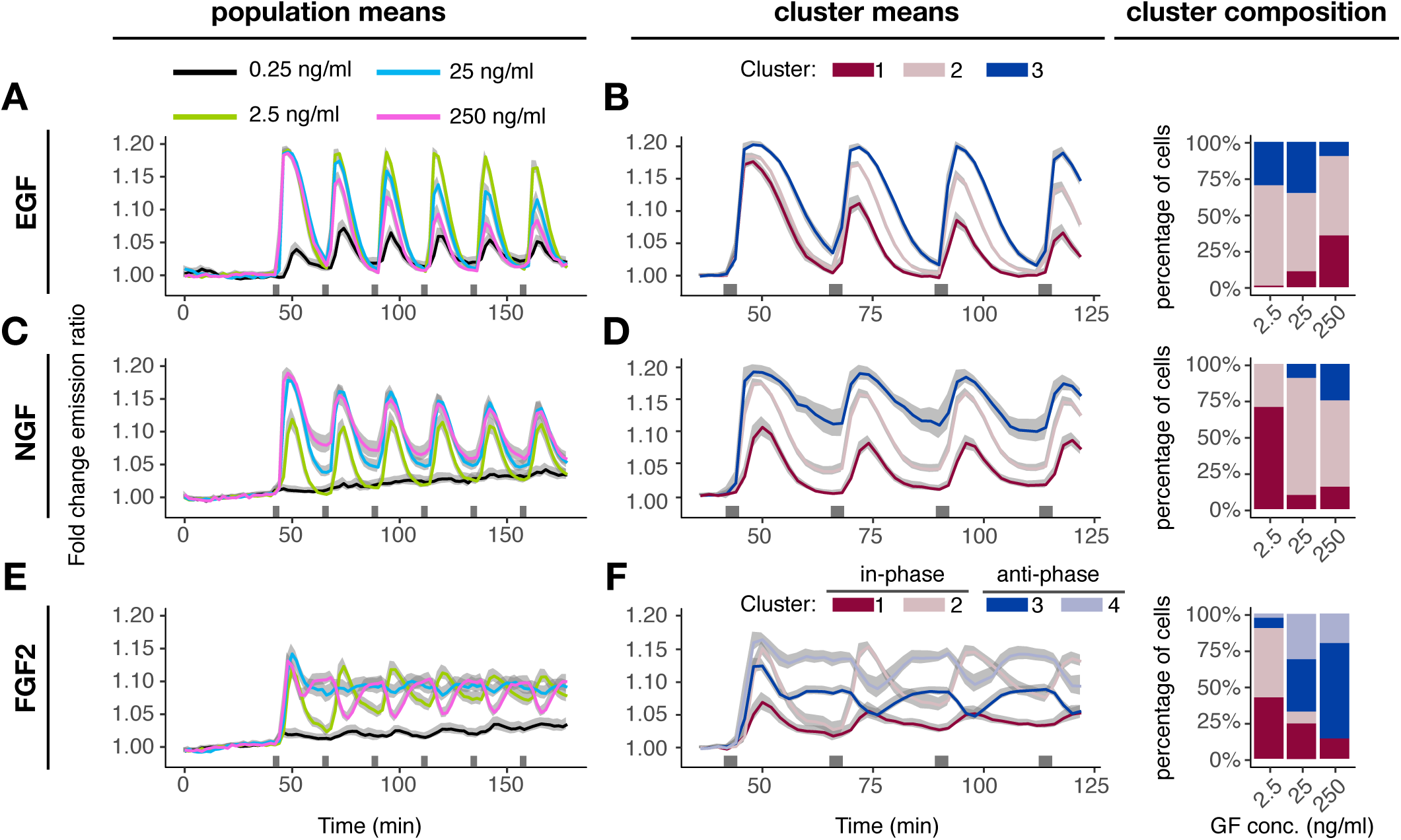
ERK activity dynamics in response to multi-pulse stimulation. (A,C,E) Population average ERK activity dynamics in response to a multi-pulse 3’-20’ EGF (A), NGF (C), FGF2 (E) stimulation. Single-cell time series were normalized to their own means before the GF stimulation, t = [0, 40]. (B,D,F) Cluster averages of ERK activity and distribution of single-cell trajectories across clusters. We performed hierarchical clustering with Manhattan distance and complete-linkage method; we cut the dendrogram at 3 (B,D) and 4 clusters (F) for visualization. Solid lines – population mean, N=[52, 91]; grey bands – 95% CI for the mean; black horizontal bars – duration of GF stimulation.

Intriguingly, varying FGF2 dosage and pulse duration again revealed more complex paTEPs than for EGF/NGF (Fig 2C). At a threshold input, we observed a new dynamic pattern, whereby an initial adaptive ERK activity peak was followed by a rebound that then decayed slowly. This dynamic pattern was visible at 25 ng/ml 10’ pulse and at 250 ng/ml 3’ and 10’ pulse. Lower or shorter FGF2 dosages induced only transient ERK activities. Clustering revealed a transient cluster as well as the characteristic ERK activity with a rebound (Fig EV3C). The latter cluster was enriched at high FGF2 inputs.

The 60’ FGF2 pulse led to even more complex paTEPs. The pattern with ERK activity rebound emerged at 2.5 – 250 ng/ml FGF2, and for these concentrations the rebound ensued only after GF was washed away. The adaptation after the initial peak was stronger at higher GF dosages. In contrast, at the lower concentration of 0.25 ng/ml FGF2, we observed sustained activation during the GF pulse and a slow decay after GF washout. Clustering of responses to 60’ pulse confirms that the lowest FGF2 dosage induces high and low sustained responses without a rebound, while higher GF concentrations result in a much stronger adaptation after the initial peak.

To probe network architectural features that might work at longer timescales and to test how the MAPK network responds to novel GF inputs before full adaptation, we subjected PC-12 cells to multiple 3’ pulses separated by 20’ pauses (Fig 3A). Multiple pulses of EGF led to transient paTEPs with 20’ timescale adaptation that were in-phase with the pulse pattern (Fig 3A). As previously shown (Ryu et al., 2015), increasing EGF concentrations correlated with increased ERK activity peak amplitude desensitization over the timescale of hours. Clustering confirmed homogeneous scTEPs across the population (Fig 3B). Multi-pulse 3’-20’ NGF also led to transient paTEPs that were in phase with the stimulation pattern (Fig 3C). Again, as previously described (Ryu *et al*, 2015), desensitization occurred at the timescale of hours, but this was not dependent on the NGF concentration as for EGF. Additionally, adaptation of individual ERK activity pulses weakened with an increased NGF dosage. Clustering indicated that this phenomenon emerged from a mix of cells with different adaptive strengths, with weakly adaptive subpopulation dominating higher NGF concentrations (Fig 3D).

In contrast, FGF2 multi-pulse datasets revealed distinct paTEPs (Fig 3E, Movie (Fig EV1)) than those seen with EGF and NGF. Consistently, with the single-pulse data, 0.25 ng/ml FGF2 pulses did not activate ERK, while 2.5 ng/ml FGF2 pulses did induce adaptive ERK activity peaks in-phase with the GF stimulation. 25 ng/ml FGF2 led to an initial peak followed by sustained ERK activity of amplitude lower than that of the initial peak. 250 ng/ml FGF2 pulses led to ERK activity peak immediately followed by short adaptation and a rebound phase, ultimately leading to sustained ERK activity. Re-triggering with the second pulse then led to immediate adaptation, followed by recovery to sustained ERK activity levels. Thus, from the 2^nd^ pulse on, ERK activity was anti-phasic with respect to the stimulation pattern. To evaluate scTEPs associated with population averages, we used hierarchical clustering with Euclidean distance to preserve information about any potential phase-shift (Fig 3F). We also omitted the 0.25 ng/ml FGF2 dataset to avoid non-responding cells. We identified four clusters, two of which resembled the in-phase ERK activity induced by the 2.5 ng/ml FGF2 (clusters 1 and 2), while the remaining two resembled the anti-phase ERK activity evoked by the 250 ng/ml FGF2 (clusters 3 and 4). Indeed, 2.5 ng/ml FGF2 consist of in-phase clusters, while 250 ng/ml FGF2 is dominated by anti-phase scTEPs. 25 ng/ml FGF2 evoked a roughly 50% population distribution of in-and anti-phase scTEPs, explaining the emergence of sustained ERK activity in population average measurements. Together, these datasets indicate the existence of different MAPK network circuitries downstream of the three GFs, with FGF2 being able to evoke more distinct signaling states than EGF/NGF (e.g. a phase shift at high FGF2 concentrations). The FGF2 concentration-dependent gradual emergence of the ERK phase shift in muti-pulse experiments parallels FGF2’s ability to gradually shift the population distribution of transient/sustained ERK states in response to sustained stimulation.

### Evaluating the role of HSPGs in the FGF2 signaling responses

HSPGs are important modulators of FGF2 signaling and enable biphasic signaling responses (Kanodia et al., 2014). To evaluate the role of HSPGs, we took advantage of the widely used chlorate (NaClO_3_) treatment to inhibit HSPG sulfation, and thus binding of FGF2 to HSPGs (Ornitz and Itoh, 2015). We benchmarked the effect of that perturbation against the FGF2-specific gradual phase-shift in ERK activity in response to increasing input in a multi-pulse experiment. We gradually inhibited HSPG sulfation with 10, 25, 50 mM NaClO_3_, and evaluated binding of a fluorescently-labeled FGF2 to (un-)perturbed cells (Fig 4A,B). We observed increasing quanta of FGF2 fluorescence retained after each pulse in unperturbed cells, likely reflecting incremental HSPG binding, as well as endocytosed material. In contrast, gradual HSPG inhibition resulted in progressive loss of FGF2 binding to the cell, with almost no remaining FGF2 at 50 mM NaClO_3_. Non-treated cells exposed to 2.5 ng/ml FGF2 pulses displayed the typical in-phase pattern. Gradual NaClO_3_-mediated HSPG inhibition led to blunting of the ERK activity amplitudes that however remained in-phase, with only low residual ERK activity at 50 mM NaClO_3_ (Fig 4C). While untreated cells exposed to 250 ng/ml FGF2 displayed the typical anti-phasic ERK activity, gradual NaClO_3_-mediated HSPG inhibition progressively rescued in-phase ERK activity, at the same time decreasing ERK activity amplitude (Fig 4D). These results show that FGF2’s ability to evoke ERK phase shifting in multi-pulse dose response experiments depends on HSPG/FGF2 interactions.

**Figure 4.**
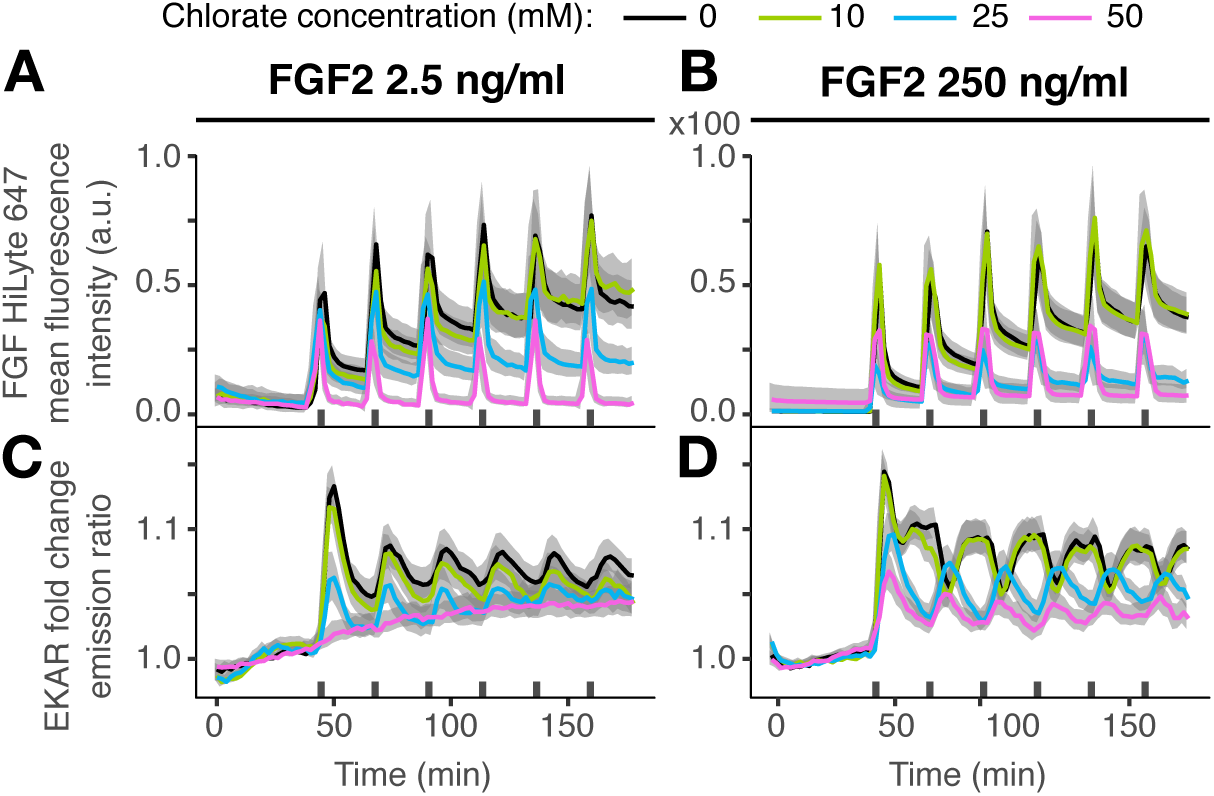
ERK activity dynamics in response to HSPG perturbation. (A,B) Population average ERK activity dynamics in response to a dose-response challenge using 10, 25, and 50 mM of NaClO3 and a multi-pulse 3’-20’ stimulation with FGF2 2.5 ng/ml (A), and 250 ng/ml (B). Top panels show a whole-cell mean fluorescence intensity of labelled FGF2 (HiLyte 647). Solid lines – population mean, N=[43, 65]; grey bands – 95% CI for the mean; black horizontal bars – duration of GF stimulation.

### Cell fate decision in response to sustained and pulsed GF stimulation

We then set out to correlate ERK activity dynamics with fate decisions. We repeated select sustained and pulsed dose response experiments and evaluated the differentiation fate by quantifying neurite outgrowth using an automated image segmentation pipeline (Fig 5A,B). 0.25 ng/ml EGF led to a relatively high level of differentiation, which is consistent with its ability to induce slowly-adapting, almost sustained ERK signaling (Figs 1D-G, EV1A). Higher EGF dosages that cause faster ERK adaptation resulted in lower cell differentiation. We observed low levels of differentiation at 0.25 ng/ml NGF, which is consistent with low amplitudes of scTEPs at this GF concentration. 2.5 – 250 ng/ml NGF led to potent differentiation, which is consistent with largely sustained scTEPs. The FGF2 induced a bi-phasic dose response in differentiation levels. At 0.25 ng/ml FGF2 we observed low levels of differentiation, which does not correlate with a mix of high and low amplitude sustained scTEPs (Fig 1D,F,G). 2.5 and 25 ng/ml FGF2, which consist of a mix of robust transient and sustained scTEPs led to relatively high differentiation, however not to the level evoked by NGF. Finally, 250 ng/ml FGF2 dominated by transient scTEPs displayed lower levels of differentiation than 2.5 and 25 ng/ml FGF2. Except for 0.25 ng/ml FGF2, a qualitative correlation exists between scTEPs and the differentiation fate.

**Figure 5.**
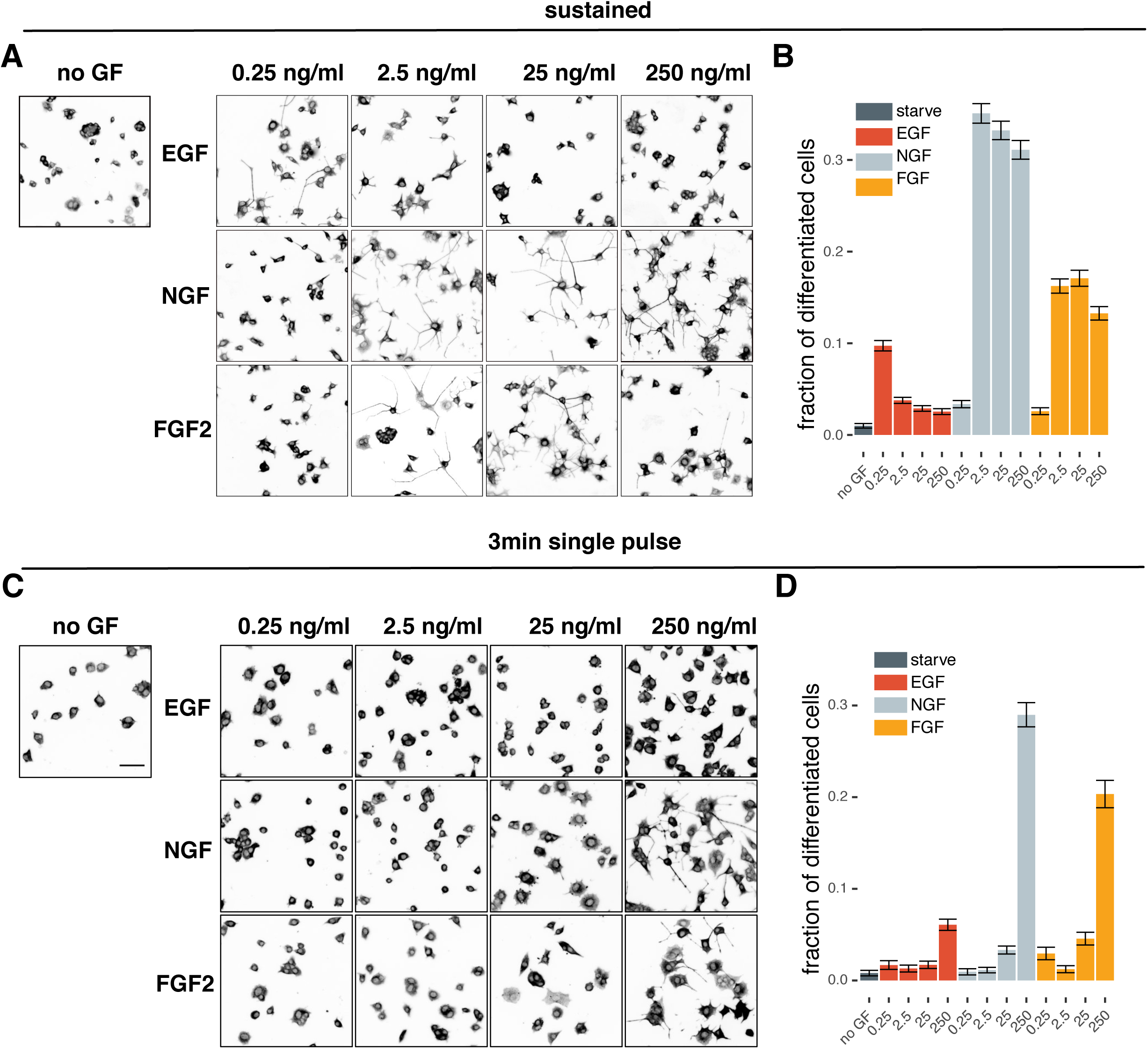
Differentiation fate analysis in response to sustained/pulsed, GF dose response stimulation. (A,C) Representative images of differentiation experiments of sustained (A) or single 3 min pulse (C) stimulation with respective GF. Cells are stained for ⟨-tubulin, and are shown in inverted black/white contrast. Scale bar = 50 ⌈m. (B,D) Fraction of differentiated cells from sustained (B) and 3 min single pulse (D) stimulation, calculated as fraction of cells with the total neurite outgrowth longer than the diameter of the cell soma using a CellProfiler-based automated image analysis routine. On average, 4000 cells per condition were included in the analysis (min = 1600, max = 12500). Error bars indicate 95% CI for the mean.

Similar to what we have shown for EGF/NGF (Ryu et al., 2015), we asked if induction of different signaling states through dynamic GF stimulation, would allow us to manipulate cell fate decisions (Fig 5C,D). We reasoned that a 3’ pulse of 250 ng/ml FGF2 that leads to an ERK activity peak followed by a sustained rebound that lasts hours (Fig 2C), would result in high differentiation. In contrast, a 3’ pulse stimulation at a lower FGF2 dosage, which does not induce ERK activity rebound should not induce differentiation. We indeed observed a correlation between this specific signaling state and differentiation. As expected, control 3’ pulsed EGF stimulation did not lead to differentiation, except for some low levels of differentiation at 250 ng/ml. We attribute this spurious differentiation to low levels of remaining EGF that cannot be completely washed out after the pulse application by pipetting, thus recapitulating the 0.25 ng/ml EGF sustained stimulation results (Fig 5A,B). The 3’ 250 ng/ml NGF pulse, also led to potent differentiation, which correlated with its ability to induce sustained ERK activity. These results indicate that sustained ERK signaling states, evoked by specific sustained or pulsed GF stimulation, correlate with the differentiation fate for all three GFs.

### Modelling the FGF2-evoked ERK activity responses

We then sought to find a minimal signaling network topology that could recapitulate the highly specific ERK signaling responses evoked by FGF2 stimulation. We reasoned that the salient features visible in dynamic ERK signaling states evoked by sustained and pulsed GF stimulation would discriminate among candidate model topologies using a Bayesian nested sampling (NS) inference method (Skilling, 2006). NS computes the posterior distribution of parameter sets for a candidate model based on experimental training datasets, and infers parameter ranges that can best reproduce the data.

We postulated a set of minimal models that aim to explain the FGF2-induced ERK activity responses. The models differed with respect to receptor interactions and intracellular signaling topologies. Three models of FGF2/HSPG/FGFR interactions (Fig 6A) were as follows. 1) Simple activation model, whereby sequential FGF2-HSPG, FGF2-HSPG-FGFR interactions ultimately lead to formation of a FGF2-HSPG-FGFR dimer and subsequent FGFR activation. The increase in FGF2 input activates FGFR until all receptors saturate (Fig 6A, right panel). 2) Competitive activation model, whereby FGF2 can also directly bind to FGFR, although this complex does not activate the receptor. Assuming that FGF2-FGFR binding occurs faster than the FGF2-HSPG binding, a biphasic activation can emerge as previously proposed (Kanodia et al., 2014) (Fig 6A right panel). Low/medium FGF2 concentrations lead to FGFR activation, while high FGF2 dosage titrates FGFR, precluding the formation of signaling-competent FGF2-HSPG-FGFR complexes. 3) Competitive joint-activation model. In contrast to the competitive activation model, the FGF2-FGFR complex is signaling-competent as previously proposed (Ornitz and Itoh, 2015). The signaling strength of FGF2-FGFR complex can be different from the FGF2-HSPG-FGFR complex, which allows residual levels of receptor activation, even at high FGF2 concentrations (Fig 6A, right panel).

**Figure 6.**
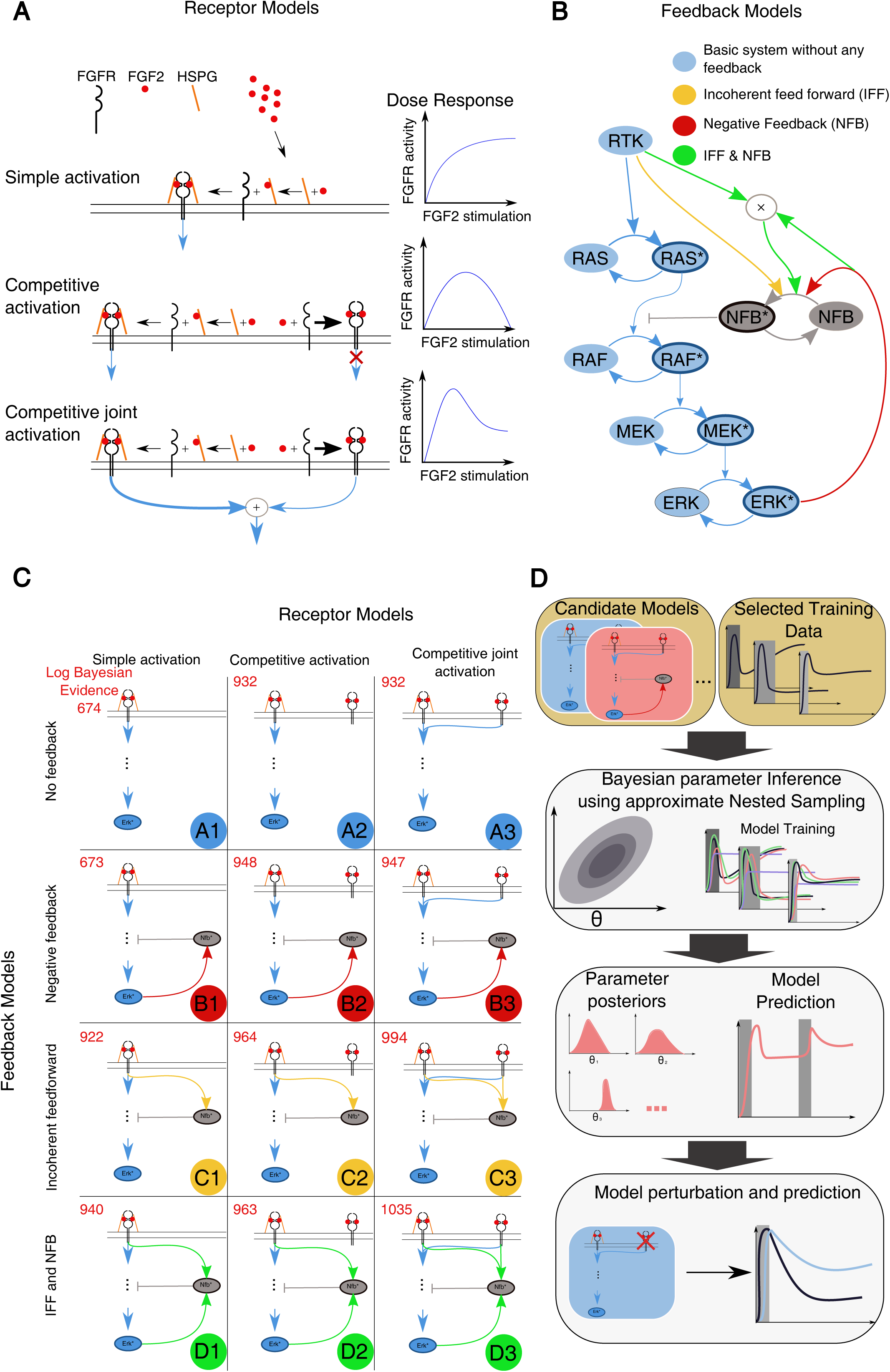
Description of candidate FGFR/FGF2 network models. (A) Illustration of the three proposed receptor activation mechanisms. Simple activation: only the FGFR-FGF2-HSGAG complex activates downstream MAPK signaling (blue arrow). Competitive activation: same as simple activation, but FGFR can bind FGF2 without HSGAG. The resulting FGFR-FGF2 complex does not activate downstream pathways. Competitive joint activation: same as competitive activation, but both FGFR-FGF2-HSGAG and FGFR-FGF2 activate downstream MAPK signaling. Right panel indicates how FGFR activity will evolve upon FGF2 stimulation. (B) Proposed MAPK network feedback models. The minimal set of MAPK signaling nodes is shown, including a negative feedback regulator protein (NFB). Asterisks denote activated forms of these nodes. Basic system: no feedback or feed-forward structure. Incoherent feed-forward: activation of the negative regulator is proportional to FGFR activation. Negative feedback: activation of the negative regulator is proportional to ERK activation. Incoherent feed-forward and negative feedback: activation of the negative regulator is proportional to the product of RTK activation and ERK activation. (C) A systematic overview of all proposed model topologies. The models are labelled A-D for different intracellular feedback structures, and 1-3 for the different receptor models. (D) An overview of our model selection procedure. We start with our set of 12 candidate models and a training set of experimental observations. For each of the candidate models, we perform Bayesian parameter inference using the approximate nested sampling algorithm. We obtain a posterior distribution of the parameters for each candidate model. We then subsequently benchmark models/associated parameter spaces for their ability to predict ERK activity dynamics for unobserved pulsed stimulation schemes. Finally, we further evaluate the ability of the candidate models to predict a biological perturbation (HSPG perturbation).

For the intracellular signaling layer, we simplified the MAPK network by taking into account the Ras GTPase, as well as the Raf/MEK/ERK kinase cascade that itself allows the production of switch-like ERK activity (Fig 6B). We considered 1) A basic model without any feedbacks. We then added three different wirings to get the following models: 2) A generic negative feedback from ERK to Raf as previously proposed (Santos et al., 2007). 3) An incoherent feedforward (IFF) system in which RTK modulates an intermediate negative regulator of Raf, termed negative feedback protein (NFB), based on our recent finding (Ryu et al., 2015). 4) A hybrid system in which the negative regulator of Raf is modulated both by an RTK-based IFF, and a negative feedback from ERK to Raf in a multiplicative way (Ryu et al., 2015). By combining the three extracellular layer receptor models with the four intracellular MAPK network feedback models, we obtained a total of 12 candidate models (Fig 6C). The models consist of 9 to 15 species and up to 39 parameters. To account for the nonlinear nature of FRET measurements, we explicitly model the ratiometric ERK activity measurement as previously described (Birtwistle et al., 2011).

To avoid overfitting, we restrained the training datasets to four experimental conditions that capture features specifically relevant to FGF2/MAPK signaling: 1. Sustained 2.5 ng/ml FGF2 that exhibits only low level of post-ERK activity peak adaptation (Figs 1D, Fig EV1C). 2. Sustained 250 ng/ml FGF2 that exhibits high level of post-ERK activity peak adaptation followed by slow recovery of ERK activity (Figs 1D, EV1C). 3. 2.5 ng/ml FGF2 multi-pulse that exhibit population-homogeneous ERK activity pulses in phase with the stimulus (Fig 3E). 4. 250 ng/ml FGF2 multi-pulse that exhibits population-homogeneous anti-phasic ERK activity pulses (Fig 3E), and thus also captures the capability of ERK activity to rebound immediately after adaptation when a high concentration FGF2 pulse is applied.

For each of the 12 candidate models, we used NS to infer the parameters that explain the dynamic ERK activity observed in four training sets. We plotted the simulations for 500 sampled parameter vectors from the posterior as well as for the best parameter set (maximum likelihood estimate), and compared it to the corresponding experimental data (Fig EV4). Visual inspection indicated that except models A1 and B1, the 10 remaining models faithfully reproduced most of the features of the training sets. The only model able to stringently explain all observations of the training dataset was the most complex model D3. The remaining models succeeded in reproducing the anti-phasic dose response for the pulse stimulation. Models A2, A3, C2, C3 and D2 failed to reproduce the initial peak, but succeeded in reproducing the ERK rebound upon sustained stimulation.

To further discriminate the models, we reasoned that we could further benchmark them against FGF2 stimulation patterns that were not used for training. With our microfluidic setup, we induced a 5’ single pulse stimulation (Appendix Fig S1A), and a more complex multi-pulse stimulation (3’ pulse/30’ pause/20’ pulse/60’ pause/5’ pulse), both at 2.5 and 250 ng/ml FGF2 (Appendix Fig S1B). The latter induces ERK responses constrained by multiple feedbacks of the FGF2/MAPK signaling within one experiment, thus resulting in dynamics not seen in our previous experiments. For each of the 10 remaining models, we sampled 500 parameter sets from each posterior and plotted the simulated ERK activity using the best-fit to the experimental dataset (Fig EV5). After visual evaluation, we excluded models that did not reproduce general trends of the experimental datasets, and we arrived at 4 models: B2, B3, C3, D3. These allow for direct binding of FGF2 to the FGFR in addition to the canonical FGFR/HSPG interactions, but display different intracellular feedbacks.

Given the importance of FGF2/HSPG interactions, we used the HSPG perturbation dataset to further discriminate the models. We set the HSPG concentration to zero and simulated the 4 remaining models using the maximum likelihood estimate of the training data (Fig 7A). The model B3 was the only one that recapitulated our experimental NaClO_3_ perturbation (Fig 7B). Thus, this model reproduced the training data, predicted unobserved ERK activity resulting from alternate dynamic stimulation, and predicted the effects of a biological perturbation (Fig 6D). Our sequential model selection procedure is depicted in Figure 6B.

**Figure 7.**
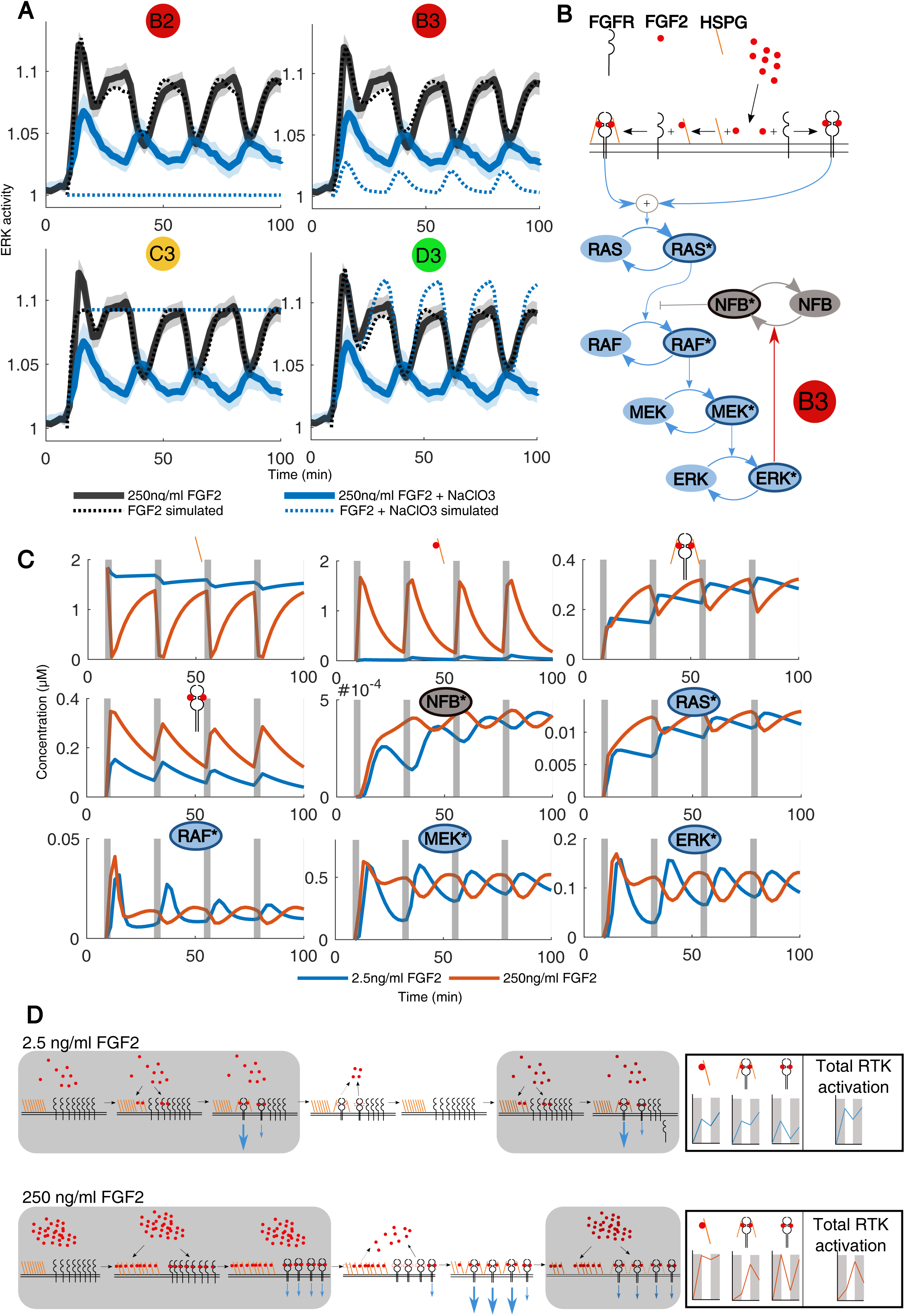
Benchmarking models against HSPG perturbation, and description of model latent states. (A) Population average ERK activity dynamics upon stimulation with 250 ng/ml FGF2 with (blue) and without NaClO3 (black) treatment (solid lines). The model simulations for the four models B2, B3, C3 and D4 are indicated with dotted lines. (B) Full illustration of the model B3, which is able to reproduce the training data, predict ERK activity dynamics upon unknown stimulation schemes, and reproduce the ERK activity dynamics upon NaClO3 treatment. (C) The latent states of the model B3 upon pulse stimulation with 2.5 ng/ml FGF2 (blue) and 250 ng/ml FGF2 (red). Different molecular species are indicated by symbols (receptor models) or signaling nodes. Asterisks denote activated forms of signaling nodes. (D) Illustration of the mechanism responsible for the in-and anti-phase ERK dynamics evoked by 2.5 and 250ng/ml FGF2. Grey boxes indicate stimulation with FGF2, blue arrows indicate signaling strengths of different FGF2 receptor complexes. Items with dotted lines indicate unbinding events. (E) Qualitative illustration of the abundance of the different receptor species, as well as total FGFR signaling activity. Grey areas indicate FGF2 pulsed stimulation.

## Discussion

An important question in the signaling field is how different RTKs, despite converging on a small number of core signaling processes (including MAPKs), can induce different fates (Lemmon and Schlessinger, 2010). An emerging concept is that the MAPK network is wired differently downstream of specific RTKs to generate distinct dynamic ERK states (Marshall, 1995; Santos et al., 2007). The latter are subsequently integrated into fate decisions through additional signal integration layers (Gillies et al., 2017; Murphy et al., 2002; Uhlitz et al., 2017). Here, we extend this notion by showing that FGF2, which has been poorly studied with respect to MAPK signaling dynamics, wires the MAPK network with a different logic, to induce distinct ERK states.

### FGF2 dose response gradually shifts the population distribution of ERK states

We report that FGF2 evokes strikingly different single-cell ERK states than EGF/NGF. Increasing FGF2 doses over 4 orders of magnitude gradually shifts the population distribution of ERK states from sustained towards adaptive response, albeit with a slow, long term rebound (Fig 1E-G). This is consistent with the previously documented population-averaged biphasic ERK activity dose responses and phenotypic outputs (Kanodia et al., 2014; Zhu et al., 2010). In contrast, increasing EGF doses trigger purely adaptive responses, and NGF produces both adaptive and sustained responses, with a steep shift towards the latter at high input. These GF identity/concentration-dependent population distributions of ERK states are quantified using cluster decomposition (Fig 1G), PCA (Fig EV2A) and a distance metric to compare the separability of time courses (Figs 1H, EV2B,C). These results indicate that a single time-lapse recording of ERK activity from the sustained FGF2 dose response challenge distinguishes low and high concentrations with a higher certainty than for EGF or NGF.

Pulsed FGF2 dose response experiments also indicated FGF2’s ability to gradually induce distinct and wider ERK state population distributions than EGF/NGF. Increasing levels of pulsed FGF2 gradually led to robust switch-like activation, strong adaptation, followed by a clear rebound leading to sustained ERK activity (Figs 2C, EV3C), with adaptation remaining for the pulse duration in the 60’ pulse experiment (Figs 2C, EV3C). In the multi-pulse dose response experiment, increasing FGF2 gradually led to a phase-shift of ERK activity patterns relative to the GF pulse (Fig 3) that depended on HSPG interactions (Fig 4), while EGF/NGF were unable to produce such a phase shift. These results again highlight FGF2’s ability to gradually shift the ERK state population distribution and thus to translate increasing GF inputs into more clearly distinct signaling states than EGF/NGF.

### Mechanistic insight into the FGF2-MAPK signaling network

To understand the network circuitry underlying FGF2-evoked ERK signaling states, we took advantage of the salient features evident in our array of sustained/pulsed experiments combined with a Bayesian inference approach. Our model selection approach consists of the following steps (Fig 6D): Formulation of minimal models that capture the relevant biology of the signaling system using *a priori* knowledge; carrying out Bayesian inference of the parameter space for each candidate model upon training on information-rich ERK states using temporal perturbations; Benchmarking model performance by predicting unknown stimulation schemes not used for training, and HSPG perturbation. We identified a simple network topology that recapitulates the ERK states observed in all these experiments. The model consists of a competitive joint activation at the receptor level (both FGF2/HSPG/FGFR and FGF2/FGFR complexes contribute to signaling), as well as a negative feedback loop from ERK to RAF (Fig 7B) – a structure recurrent in many MAPK networks (Birtwistle and Kolch, 2011; Santos et al., 2007). Thus, an interplay between a ligand, a co-receptor, and a receptor that leads to competitive activation of two signaling-competent complexes, coupled to a simple intracellular negative feedback can recapitulate all observed FGF2-dependent ERK states.

To provide intuition about the interplay of the different molecular species involved in the network, we plotted their latent states. For the sake of simplicity, we focus on the robust, population-homogeneous, in-and anti-phase ERK states evoked by multi-pulse stimulation by 2.5 and 250 ng/ml FGF2 (Fig 7C). Schemes of the receptor interactions/activities are also provided for clarity (Fig 7D,E). At 2.5 ng/ml FGF2, FGF2 binds to FGFR and HSPG, increasing the HSPG+FGF2, FGFR+HSPG+FGF2 concentration and leading to strong FGFR activity, as well as increasing the concentration of FGFR+FGF2 which leads to low FGFR activity (Fig 6D, right panel). Upon FGF2 washout, FGF2 unbinds from the 3 receptor complex species. Thus, the FGF2 pulse initially leads to combined high FGFR activity, that subsequently adapts to some extent upon FGF2 washout, translating into a RAF/MEK/ERK activation/deactivation cycle with robust adaptation due to negative feedback. During subsequent FGF2 pulses, further increase in global FGFR activity is observed, explaining the in-phase ERK activation pattern. At 250 ng/ml FGF2, much more HSPGs will be FGF2-bound, mostly leading to the formation of HSPG+FGF2 and FGFR+FGF2 species at the cost of FGFR+HSPG+FGF2 complex formation. During the 1^st^ pulse, global FGFR activation thus results from formation of some FGFR+HSPG+FGF2, but to a higher extent of FGFR+FGF2 complexes. Accordingly, RAF, MEK and ERK get activated very similarly to the 2.5 ng/ml FGF2 pulse due to the switch-like activation enabled by the tripartite structure of the MAPK network. However, after the pulse FGF2 unbinds from FGFR, leaving the still abundant HSPG+FGF2 species to engage into FGFR+HSPG+FGF2 complexes, with strong FGFR activity. After adaptation due to strong negative feedback, this leads to a delayed increase in RAS, RAF, MEK and ERK activation. At the 2^nd^ pulse, high FGF2 levels saturate HSPGs and enhance FGFR+FGF2 at the expense of FGFR+HSPG+FGF2 complexes, leading to a decrease in overall FGFR activity and to subsequent RAS, RAF, MEK and ERK inactivation. An important feature captured by the model, which is already mentioned above, is that the HSPG+FGFR+FGF2 complex signals much more strongly than just FGFR+FGF2. This can be observed because Ras activity mostly follows the profile of HSPG+FGFR+FGF2, rather than that of FGFR+FGF2 species formation. Indeed, the model prediction that the HSPG+FGFR+FGF2 complex signals more strongly than the FGFR+FGF2 complex is expected since the latter has been proposed to be the canonical, stable signaling unit (Ornitz, 2000). These results indicate how the ERK activity phase shift emerges with increasing FGF2 concentrations through modulations of the abundance of different receptor species with differential signaling ability. We propose that the same receptor competition mechanism enables the gradual shift of the population distribution of transient/sustained ERK activity states in the sustained stimulation dose response.

### Distinct characteristics of specific RTK-MAPK signaling systems

We showed that different RTKs and their cognate GFs produce distinct scTEPs population distributions. Our results extend the notion that at least some of this specificity is encoded in the MAPK network. We demonstrated that the EGF, NGF, and FGF2 signaling pathways can sense almost 4 orders of magnitude of GF concentrations and translate them into robust signaling states that are biologically relevant for fate decisions.

The EGF/NGF systems interpret graded inputs into a variety of signaling states, thanks to the modulation of negative/positive feedbacks by their respective RTKs (Ryu et al., 2015). At all concentrations (0.25 – 250 ng/ml), sustained EGF induces switch-like, adaptive responses with a robust high-amplitude initial peak that narrows as the GF dosage increases (Fig 1D-G). Hence, gradual EGF increase leads to a stronger negative feedback and thus faster robust adaptation, which can protect the RTK signaling against stochastic effects due to heterogeneous expression of signaling components (Birtwistle and Kolch, 2011). Low EGF input produces slow adaptive, almost sustained, ERK activity (Figs 1D-G, EV1) sufficient to differentiate a portion of cell population (Fig 5A,B). High EGF levels lead to low differentiation due to fast adaptation of ERK activity. Thus, an EGF dose response can already produce sufficiently distinct signaling states to induce different fates. In marked contrast, low sustained NGF input induces only low-amplitude responses, but at high NGF input the responses become sustained and saturate above 2.5 ng/ml NGF (Fig 1D). Pulsed stimulation indicates a mix of adaptive/sustained ERK activity responses (Figs 2B, EV3B), with the population distribution of the latter increasing at high NGF input. This again documents our previous finding of RTK-modulated positive feedback: increased NGF input leads to stronger positive feedback, consequently increasing the fraction of cells with sustained ERK activation (Ferrell and Machleder, 1998). However, saturation of ERK activity limits the gradual increase of the portion of cells with sustained signaling (Fig 1D-G). The NGF-TrkA-MAPK network varies the distribution of adaptive and sustained responses, and thus can shift the population distribution of proliferation/differentiation fates in response to increasing input, as previously proposed (Chen et al., 2012).

In contrast to the EGF/NGF systems, the FGF2-FGFR-MAPK network interprets the input concentration using a system that comprises competing receptor complexes in the extracellular space and a simple intracellular network with a negative feedback. The robustness emerging from this negative feedback might allow FGF2 to reliably integrate a large range of FGF2 concentrations into the gradually changing mix of transient/sustained ERK states. The function of this network might be to enable reliable signal transmission of fate decisions during developmental FGF morphogen gradient interpretation (e.g. FGF2 input might gradually vary the relative abundance of transient/sustained ERK states, evoking different fates along a morphogen gradient) (Ornitz and Itoh, 2015). Our work provides novel insights about how three distinct RTKs wire the MAPK network to differentially fine-tune the population distribution of dynamic ERK signaling states and fate decisions.

## Materials and Methods

### Cell culture

PC-12 cells stably expressing the EKAR2G1 construct (described earlier Ryu et al., 2015) and PC-12 Neuroscreen-1(NS-1, gift from Tobias Meyer) where cultured using low glucose DMEM (Sigma) supplemented with 10% horse serum (HS, Sigma), 5% Fetal Bovine Serum (FBS, Sigma) and 1% Penicillin/Streptomycin. Cells were cultured on plastic tissue culture dishes (TPP) coated with 50ug/ml collagen from bovine skin (Sigma). Cells were passaged at 70% confluence by detaching cells using a cell scraper (Fisher).

### Microfluidic device fabrication and preparation

Microfluidic device preparation was performed as described previously (Ryu *et al*, 2015). In short Polydimehylsiloxane (PDMS) polymer (Dow Corning) was mixed with the catalyzer in 10:1 ratio in a plastic beaker. A first layer of 4-5g was poured on the master then degased in a dessicator before solidifying at 80°C for 1h. 8-well reservoir strips (Evergreen) were divided in 2 and then glued on the first layer using PDMS and solidifying at 80°C for 30 min. Finally, the second layer of 15-20g of PDMS is used to finalize the device. The PDMS replica was the cut and punched at the appropriate inlets and outlets. Plasma treatment was used to bond the PDMS replica to the 50×70mm coverslip (Matsunami, Japan) to allow proper sealing that resists the high-pressure applied during the experiments. To enhance the bonding strength, the device was heated for 15 min in an 80°C dry oven. After bonding, the device was immediately filled by adding 200 ul of 50ug/ml collagen solution in PBS to each outlet reservoir and put at 37°C. To increase coating efficiency in the device 10ul of the collagen solution was aspirated 3x from the cell seeding port after 1 h each before seeding cells.

PC12/EKAR2G cell suspensions were prepared at a concentration of 10^6^ cells/ml. 50 μl of this cell suspension was added in the outlet and aspirated with a pipette from the cell reservoir inlet port. After a 10′ incubation, residual cells in the outlet were removed by aspiration and 250ul of DMEM supplemented with 10% HS, 5%FBS and 1% Penicillin/Streptomycin was added to the outlet reservoir. The inlet reservoirs were filled with 150ul of starving medium (pure low glucose DMEM). Prior to experiments, outlet reservoirs were emptied and inlet reservoir were filled with 200 ul of fresh starving medium (inlet1) and starving medium with appropriate GF/NaClO3 concentration and added dextran-Alexa 546 (4nM final concentration) (inlet2).

### Live cell imaging

All FRET ratio-imaging experiments were performed on an epifluorescence Eclipse Ti inverted fluorescence microscope (Nikon) with a Plan Apo air 20× (NA 0.75) objective controlled by NIS-Elements (Nikon). Laser-based autofocus was used throughout the experiments. Image acquisition was performed with an Andor Zyla 4.2 plus camera at a 16-bit depth. Donor, FRET, and red channel images (to visualize an Alexa546-dextran that indicates GF exposure) were acquired sequentially using filter wheels. The following excitation, dichroic mirrors, and emission filters (Chroma) were used: donor channel: 430/24×, Q465LP, 480/40 m; FRET channel: 430/24×, Q465LP, 535/30 m; and red channel: ET550/15, 89000bs, 605/50 m (for dextran imaging). Standard exposure settings were used throughout the experiments. 440-nm (donor and FRET channel excitation) and 565-nm (red dextran) LED lamps were used as light sources (Lumencor Spectra X light engine), with 3% (440 nm) and 5% (565 nm) of lamp power. Acquisition times were 30 ms for donor channel and 30 ms for FRET at binning 2 × 2 and 100ms 8×8 binning for the red channel. Cells were imaged in DMEM with 1,000 mg/ml glucose, and penicillin/streptomycin, at 37°C. The microfluidic device was mounted on the microscope stage and was connected by the tubing to a cellASIC ONIX (merck millipore) pump.

### NS-1 differentiation experiments

8000 cells per well (96 well plates BD Bioscience) were seeded in starving medium consisting of low glucoses DMEM supplemented with 0.2% HS and 1% Penicillin/Streptomycin. Cells were starved for 24 hours before adding 200ul of the appropriate GF. For 10 min and 3min experiments, wells were carefully washed once with 200 μl starving medium to dilute residual GF. After 2d cells were fixed using 4% PFA at 37°C for 10min and subsequently washed twice with PBS. Cells were permeabilized in 0.1% Triton X 100 in PBS for 10’ and blocked in PBS with 1% BSA and 22mg/ml glycin for 30’. Cells were then incubated with an anti-alpha-tubulin (Sigma T9026, 1:1000) antibody at 4°C overnight, washed 3 x for 10’ with PBS, and incubated with an Alexa-546 anti-mouse secondary antibody (1:1,000) for 3 h. Samples were washed with PBS with DAPI (1:1000) for 5′ and then washed 2x 10’ with PBS before imaging.

### Image Analysis

For the segmentation, tracking, and ratio calculation in time-lapse experiments we used CellProfiler. First, FRET and Donor channels were corrected for uneven illumination using the *CorrectIlluminationCalculate* and *CorrectIlluminationApply* modules using the *Background* setting. Cells were then segmented using the *IdentifyPrimaryObjects* module. As there is no nuclear marker for segmentation, we excluded clumps of cells using stringent size exclusion in this module. We tracked objects using the *TrackObjects* module and calculated the ratio image using the *ImageMath* where the FRET image is divided by the Donor image. Using the *MeasureObjectIntensity* mean intensity of the newly created ratio image was measured for every tracked object as well as mean intensity of labeled FGF HyLight 647 for experiment using it. In addition, total image intensity of the red dextrane Alexa 546 channel was measured to follow GF exposure. Measurements were exported as csv files and quantified with R scripts.

We used a different CellProfiler pipeline to analyze differentiation experiments. First DAPI channel was segmented using *IdentifyPrimaryObjects* to detect nuclei. Using the *IdentifySecondaryObjects* module, cells including their neurites were segmented using the nuclei objects as a seed and the tubulin stain as the image. These objects were then skeletonized using the *ExpandOrShrinkObjects* module. To obtain the soma a series of morphological operations were applied (4x erode followed by 4x dilation) to the tubulin images using the “Morph” module, then resulting images were segmented again using *IdentifySecondaryObjects*. Total neurite length per cells was then measured using *MeasureNeurons* module and data was exported to csv files.

### Quantification and statistical analysis

#### Clustering

We used R software to analyze and cluster time series. The amplitude of each trajectory was first normalized to its own mean before GF stimulation, i.e. *t* ∈ [0,40] for Figures 1C,D, 2A-C, 3A,C,E, 4A,B, or *t* ∈ [36,40] for Figures 1E, 3B,D,F.

For clustering of sustained and single-pulse GF stimulations we used dynamic time warping distance from *dtw* R package. The subsequent hierarchical clustering was performed using standard R functions *dist* and *hclust*.

#### PC analysis

We use standard R function *prcomp* for principal component analysis (PCA). For the decomposition, we use pooled data for all GFs (EGF, NGF, FGF2) and their concentrations (0.25 – 250 ng/ml) from Figure 1E (main text). After the decomposition we add negative control dataset (no GF) by rotating it to the new PC basis.

#### Population distance

With this novel approach we set out to quantify the separation between two populations of single-cell time series as shown in Figure S2B. The two populations may correspond to single-cell dynamic ERK responses to two treatments with different GFs or different GF concentrations.

In the first step, at every measured time point we calculate distance between two distributions of a measured quantity (middle panel of Figure S2B). We use Jeffries-Matusita distance (dJM), which for two normalized histograms *p* and *q*, with ∑_*i*_*p*_*i*_= 1, reads:

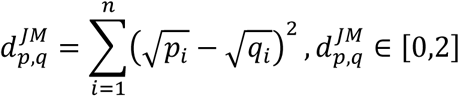

Where, *n* is the number of histogram bins. The J-M distance is bound and equals 0 for two identical distributions, and 2 for two entirely disjointed distributions regardless of how far apart they are. As a result, dJM overemphasizes small, and suppresses high separability values.

In the second step, we calculate the fraction of area under dJM curve over time, relative to maximum AUC of dJM. Therefore, the population distance between two sets of single-cell timeseries, a and b, reads:

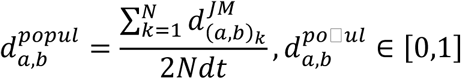

Where, *N* is the number of time points, *dt* is time interval of data, index *k* indicate a time point at which dJM is calculated.

### Mechanistic modelling

Table 1 shows all modeled species, their notation used for the equation as well as the initial values. The initial values for the FGF2 receptor, the negative feedback species as well as HSPG are inferred from the data and are modeled through the indicated parameters. The other initial values were taken from http://bionumbers.hms.harvard.edu.

The model equations for the basic model are shown in Table 2. The phosphorylation events are modeled with Michaelis-Menten kinetics. The negative feedback is modeled through the modeling species *NFB* and its “active” version *NFB*^*^which affects the phosphorylation rate of *RAF* in a hill – type manner, with hill coefficient *h*_*nfb*_. The receptor models are shown in Table 3. Since the activation happens through direct binding of FGF2 to HSPG and then to FGFR, these reactions have linear propensities. Even though the activation of FGFR requires dimerization (and possibly multimerization) we modelled it linearly to avoid making the model unnecessarily complicated.

The negative feedback is regulation depends on the model (Equations in Table 4). For the incoherent feed forward models (C1, C2, C3), it is only regulated by the membrane species *HFR* and *FGFR*^*^; for the negative feedback models (B1, B2, B3), it is only regulated through *ERK*^*^; and for the models (D1, D2, D3), it is regulated through both the receptor species *HFR* and *FGFR*^*^as well as *ERK*^*^. Since the NFB does not correspond directly to any biological species, we modeled its activation as hill dynamics with inferred hill coefficients *h*_*hfr*_and *h*_*fgfr*_.

SBML file deposited in https://www.ebi.ac.uk/biomodels/

### Parameter estimation

All parameters are listed in Table 5. For the parameter estimation we used a custom implementation of a Nested Sampling (NS) method based on (Skilling, 2006). Given the experimental data, NS approximates the posterior (being the Bayesian evidence and being the likelihood) by drawing samples of parameter vectors from the prior, along with weights that are used to recover the posterior distribution and the Bayesian evidence. NS explores areas of the prior constrained to higher regions of likelihood corresponding to an increasing sequence of thresholds. For a thorough discussion on the NS approach see for instance (Skilling, 2006). For a termination criteria, we follow (Skilling, 2006) and ran the NS algorithm until the final prior volume multiplied by the highest likelihood in that volume is smaller than 0.001 times the current evidence estimate and the difference between the highest log-likelihood and the lowest log-likelihood in the current “live” set is less than 2. The NS algorithm was run in parallel on 48 cores and the sampling from the constrained prior in each iteration was by performing density estimation on the current “live” points and using rejection sampling to sample from the prior on the support of the constrained prior. The priors for all parameters were chosen to be log-uniform.

## Data availability

Data files are available at: https://data.mendeley.com/datasets/mv3md5ty5w/draft?a=c830b09c-fefc-4cd1-bffb-38aecc7e979f

## Acknowledgements

This work has been supported by grants from the Swiss National Science Foundation to O.P., from the Korean-Swiss Science and Technology Programme to O.P. and N.L.J., from the Basic Science Research Program through the National Research Foundation of Korea (NRF) funded by the Ministry of Science and Technology (NRF 2018R1A2A1A05019550) to N.L.J, and from the European Union’s Horizon 2020 research and innovation programme under grant agreement N° 730964 (TRANSVAC project) to M.K. We thank Tobias Meyer for the PC-12 NS-1 cell line.

## Author Contributions

O.P, Y.B., M.D, H.R. conceived the study. Y.B. and H.R. performed experiments. Y.B., M.D. and M.A.J analyzed data. J.M. and M.K. performed mathematical modelling. H.R. and N.L.J set up the microfluidic platform. O.P. supervised the work. O.P., M.D., J.M., Y.B. wrote the paper.

## Expanded View Figure legends

**Figure EV1.**
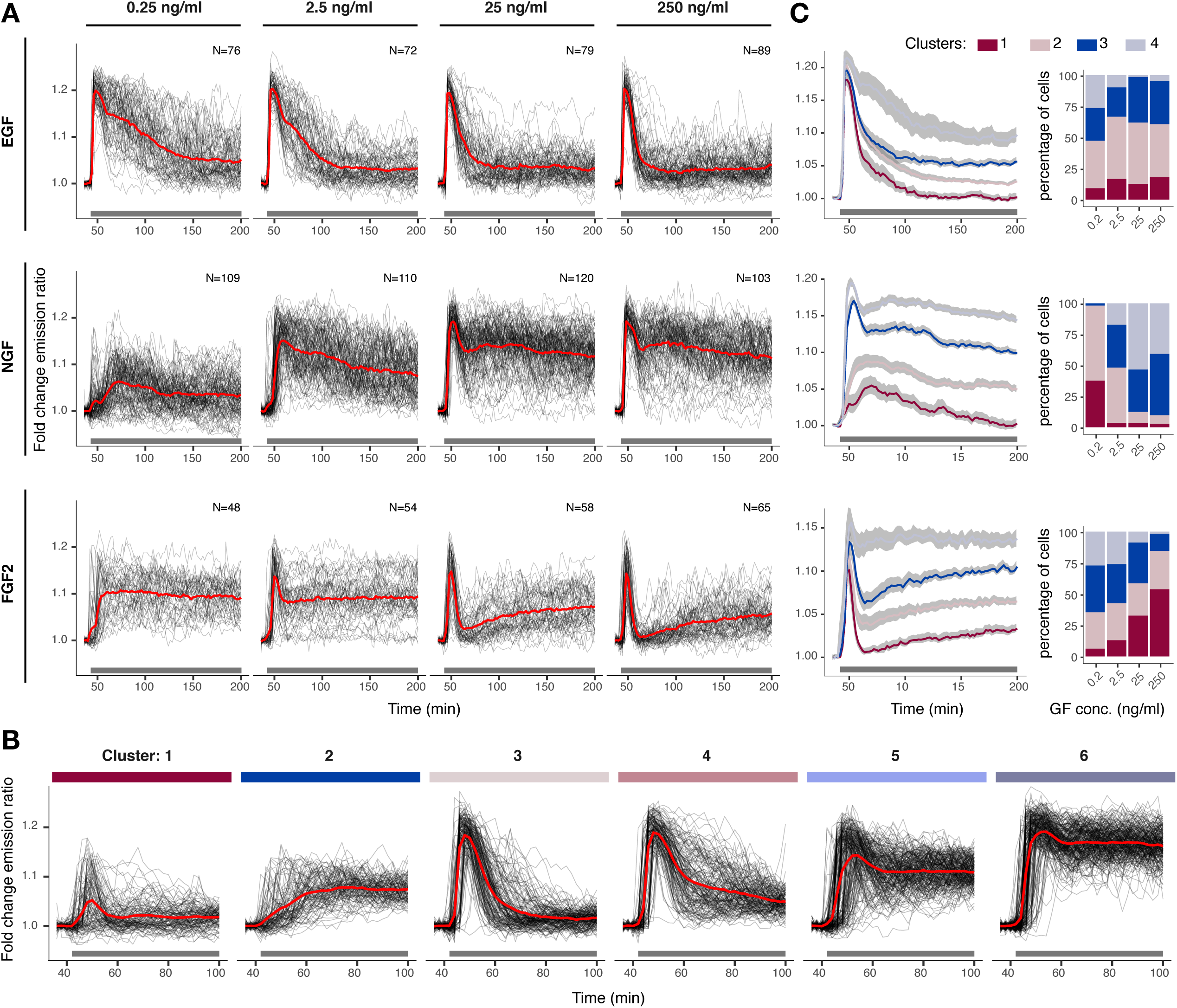
Raw single-cell ERK activity trajectories of the EGF, NGF, FGF dose response. (A) ERK activity dynamics in response to stimulation with a dose-response challenge using 0.25, 2.5, 25, 250 ng/ml EGF, NGF, FGF2. Single-cell time series were normalized to their own means before GF stimulation, t = [36, 40] min. Red curve indicates the population mean; black horizontal bar indicates GF stimulation. N=[48, 120] cells per GF concentration. ERK dynamics measured at 2’ intervals. (B) Single-cell ERK trajectories within 6 clusters identified in Figure 1F. We used hierarchical clustering with Dynamic Time Warping (DTW) distance and Ward’s minimum variance linkage method for building dendrogram, which was then cut to distinguish 6 clusters. (C) Hierarchical clustering of individual GF dose-response challenges from (A) with the same approach as (B) but for a longer interval (t = [36, 200] min). Left column indicates cluster means and 95% CI for the mean, right column is the distribution of single-cell trajectories across 4 clusters.

**Figure EV2.**
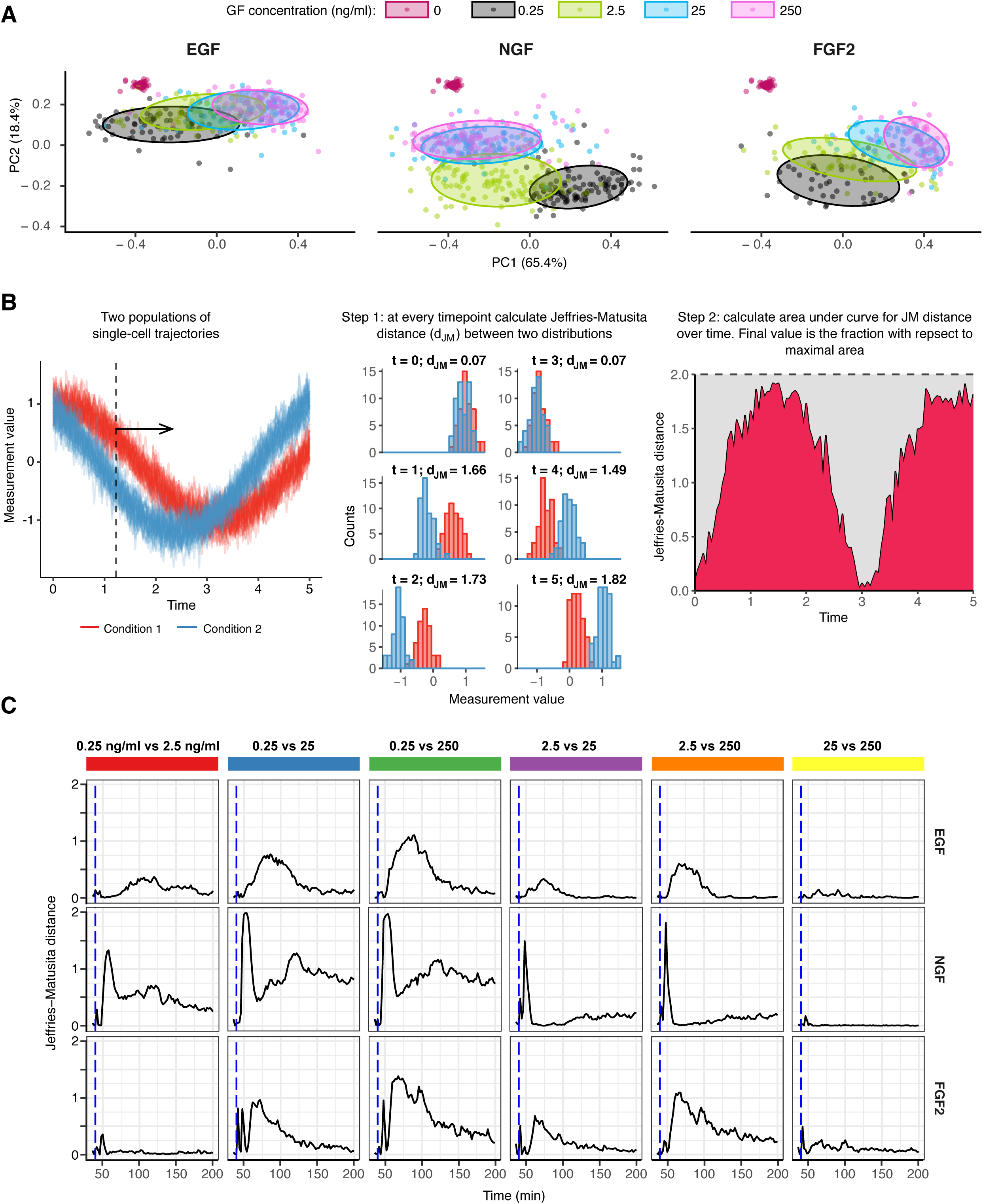
Separability of ERK states in the EGF, NGF, FGF2 dose response challenge. (A) Principle component analysis (PCA) of a pooled dataset from Figure 1E. First two components account for 85% of the overall variability. Here, we use the same data trimmed to t=[36,100] min as in clustering in Fig. 1E. Note that in addition to GF responses, responses of untreated cells are also shown, although they were not used in the decomposition (red points). (B) Schematic of distance calculation between two populations of single-cell timeseries. Two synthetic populations of noisy time series data consist of 100 trajectories each, with each trajectory spanning t=[0, 5] T with 0.1 T interval. Step 1: at every measured time point, calculate Jeffries-Matusita distance (d_JM_) between two distributions of a measured quantity. Step 2: calculate area under curve (AUC) of d_JM_ and express it as fraction of the maximum AUC of d_JM_, which is 2*dxN*, where *dx* – interval length, *N* – number of measured time points. The normalized AUC of d_JM_ is used to construct dendrogram in Fig. 1H. (C) d_JM_ over time for each GF concentration pair (0.25, 2.5, 25, 250 ng/ml) for EGF, NGF, and FGF2 stimulation. Here, we use the same data trimmed to t=[36,200] min as in Figure EV1A.

**Figure EV3.**
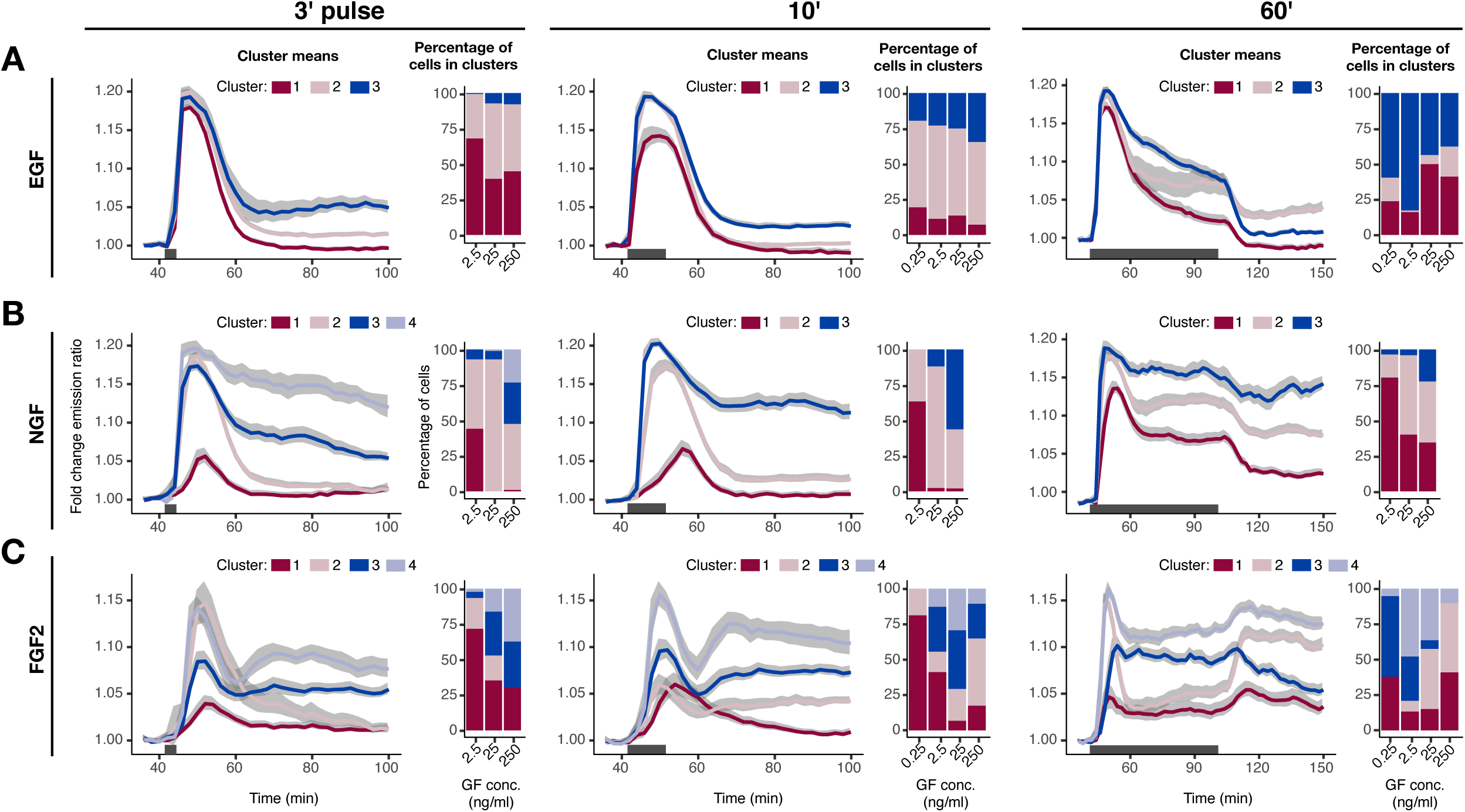
Clustering of dynamic ERK activity signaling states in response to single-pulse GF stimulation regimes. (A-C) Hierarchical clustering of dynamic ERK responses to a single 3’, 10’, and 60’ pulse of EGF (A), NGF (B), and FGF2 (C). Clustering approach was the same as in Figure EV1. We cut the dendrogram at 3 or 4 clusters for ease of visualization.

**Figure EV4.**
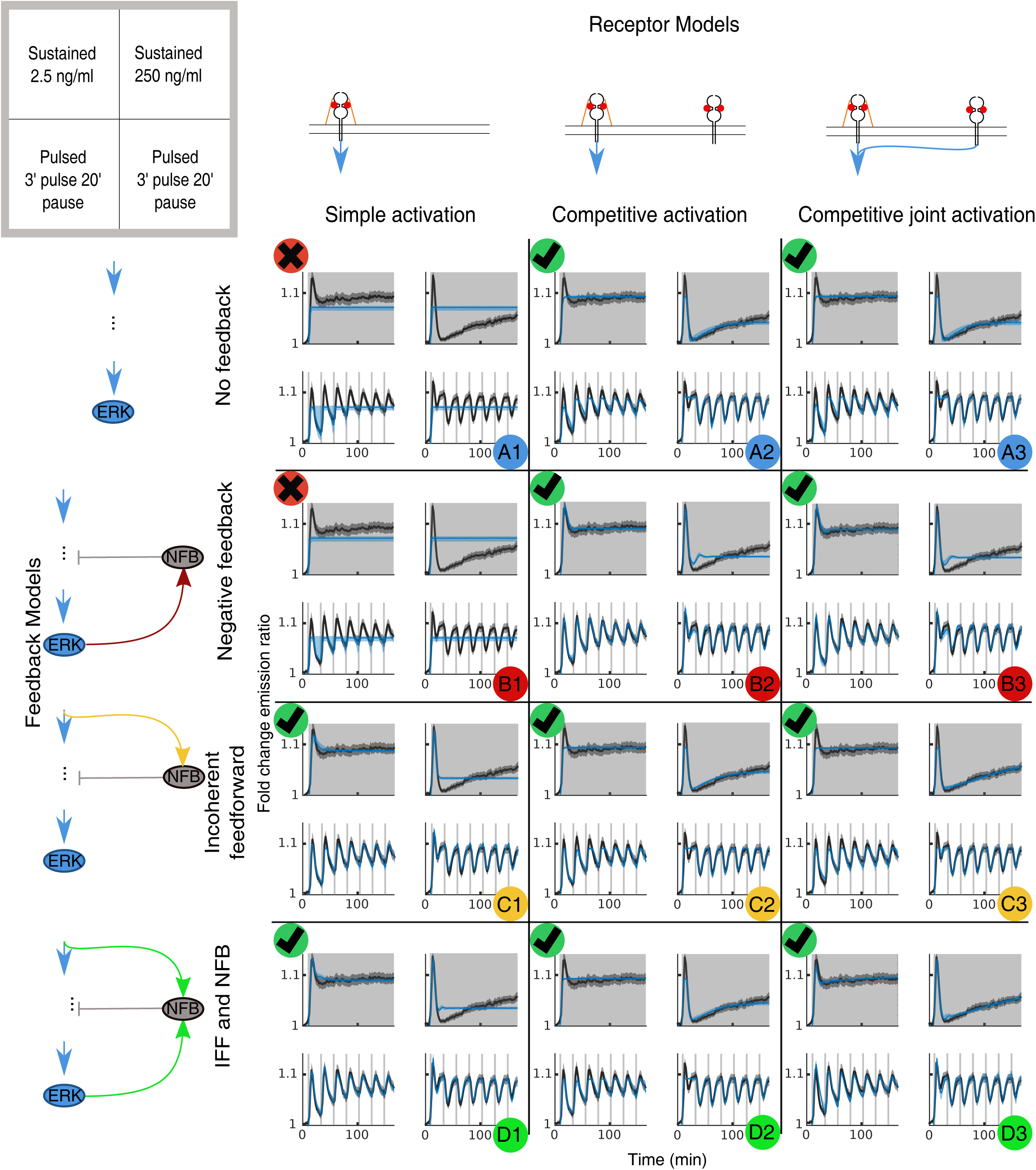
Candidate model training. Model training for all proposed candidate models. Each model was trained on ERK activity population averages of the following experimental datasets (as shown in the upper left key): 2.5ng/ml sustained FGF2 stimulation (top left), 250ng/ml sustained FGF2 stimulation (top right), 2.5ng/ml pulse stimulation (bottom left) and 250ng/ml pulse stimulation (bottom right). Experimental ERK activity population averages: black lines – 95% CI (grey shaded area). GF stimulation: light-grey vertical areas. Best fit for of each model (maximum likelihood of training): blue lines. Envelope of the simulations based on parameters sampled from the posterior distribution: light-blue areas. Models A1 and B1 fail to capture the responses and are discarded (black crosses). The remaining models are considered for further analysis.

**Figure EV5.**
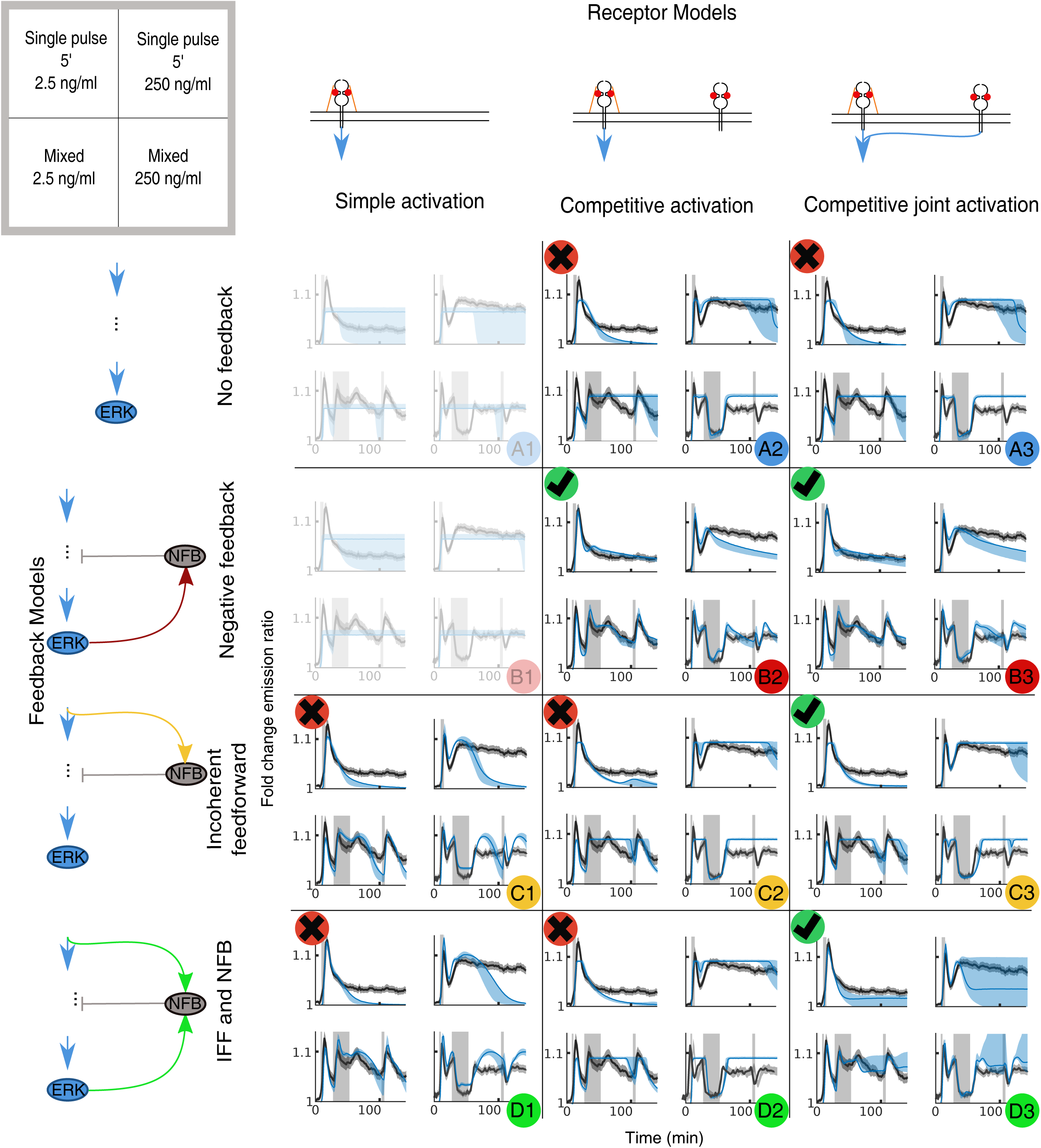
Candidate model prediction. Each model was simulated and compared with the ERK activity population averages of the following experimental datasets (as shown in the upper left key): 2.5 ng/ml FGF2 single 5’ pulse (top left), 250 ng/ml FGF2 single 5’ pulse (top right), 2.5 ng/ml FGF2 mixed pulse (bottom left), 250 ng/ml FGF2 mixed pulse (bottom right). Experimental ERK activity population averages: black lines – 95% CI (grey shaded area). GF stimulation: light-grey vertical areas. Best prediction for of each model (maximum likelihood of training): blue lines. Envelope of the simulations based on parameters sampled from the posterior distribution: light-blue areas.

